# Augmented Intelligence with Natural Language Processing Applied to Electronic Health Records is Useful for Identifying Patients with Non-Alcoholic Fatty Liver Disease at Risk for Disease Progression

**DOI:** 10.1101/518217

**Authors:** Tielman T. Van Vleck, Lili Chan, Steven G. Coca, Catherine K. Craven, Ron Do, Stephen B. Ellis, Joseph L. Kannry, Ruth J.F. Loos, Peter A. Bonis, Judy Cho, Girish N. Nadkarni

**Author notes:** Equal Contribution (Judy Cho, Girish N. Nadkarni). **Funding Support:** NIH/NIDDK. **Correspondence:** Tielman T. Van Vleck, Ph.D. or Girish N. Nadkarni, MD, MPH, CPH, Icahn School of Medicine at Mount Sinai, One Gustave L. Levy Place, Box 1243, New York, NY 10029, Email Address or.

## Abstract

**Objective:** Electronic health record (EHR) systems contain structured data and unstructured documentation. Clinical insights can be derived from analyzing both but optimal methods for this have not been studied extensively. We compared various approaches to analyzing EHR data for non-alcoholic fatty liver disease (NAFLD).

**Materials and Methods:** We compared analysis of structured and unstructured EHR data using natural language processing (NLP), free-text search, and diagnostic codes against expert adjudication as the reference standard.

**Results:** Out of 38,575 patients, we identified 2,281 patients with NAFLD. From the remainder, 10,653 patients with similar data density were selected as a control group. NLP was more sensitive than ICD and text search (NLP 0.93 vs. ICD 0.28 vs. text search 0.81) with higher a F2 score (NLP 0.92 vs. ICD 0.34 vs. text search 0.81). 619 patients had suspected NAFLD documented in radiology notes not acknowledged in other forms of clinical documentation. Of these, 232 (37.5%) were found to have more advanced liver disease after a median of 1,057 days.

**Discussion:** NLP-based approaches have superior accuracy in identifying NAFLD within the EHR compared to ICD/text search-based approaches. Suspected NAFLD on imaging is often not acknowledged in subsequent clinical documentation. Many such patients are later found to have more advanced liver disease.

**Conclusion:** For identification of NAFLD, NLP performed better than alternative selection modalities and facilitated follow-on analysis of information flow. If accuracy can be proven to persist across clinical domains, NLP can identify patient phenotypes for biomedical research in an accurate and high-throughput manner.

## INTRODUCTION

Non-alcoholic fatty liver disease (NAFLD) represents a spectrum of liver diseases characterized histologically by macrovesicular fat and ranging in severity from nonalcoholic fatty liver (NAFL) to non-alcoholic steatohepatitis (NASH).[1, 2] A subset of patients with NAFLD progresses to cirrhosis and has an increased risk of hepatocellular carcinoma and liver-related mortality.[1] NAFLD is emerging as one of the most common causes of liver failure in the United States.[3]

Multiple professional societies have published guidelines for the diagnosis and management of patients with NAFLD.[4] Clinical evaluation is warranted in patients suspected of having NAFLD including those with known risk factors (such as type II diabetes) and compatible imaging findings such as fatty liver seen on ultrasound or cross-sectional imaging.[5] The extent to which such patients are recognized in clinical practice has not been studied extensively although the available data suggest that under-diagnosis is common.[4]

Electronic health records (EHR) systems offer the potential to identify patients with NAFLD in whom the diagnosis has not been recognized or pursued. A challenge in identification of such patients is that relevant information is frequently documented in unstructured data sources such as clinic notes, discharge summaries, and radiology reports[6] in contrast to structured data, which are represented by standardized terminology such as the International Classification of Disease-Clinical Modification (ICD-CM) codes. Unlike structured data, where a concept is represented as a discrete value, an understanding of unstructured data is limited by the ambiguity of natural language. A human, for example, can easily recognize whether the expression “Paris Hilton” refers to a hotel in Paris or the celebrity based on context. Such contextual understanding has proven more challenging for computers. However, advances in augmented intelligence approaches, such as natural language processing (NLP), have shown increasing promise in harvesting rich sources of unstructured data. These approaches have been used successfully for biomedical research such as accurate phenotyping of complex diseases and for clinical tasks including identification of patients with NAFLD.[7–8]

In this study, we compare various approaches for identifying patients with NAFLD based on structured and unstructured information contained in an EHR. We then attempt to determine the proportion of patients with fatty liver documented in radiology reports in which the possible diagnosis of NAFLD was also documented in a progress note from a healthcare provider and estimated the proportion of patients with possible NAFLD who progressed to cirrhosis/liver failure. It has been shown that patient errors can be detected by accurately mining the patient record.[9] In an attempt to assess the model’s ability to identify communication breakdowns at the point of care, we also examine patients where NAFLD was identified in radiology notes but was not ever referenced in progress note and the patient still progressed to NASH or cirrhosis.

## METHODS

### Study setting and population

The study was conducted at the Icahn School of Medicine at Mount Sinai and used the data resources of the BioM? Biobank at the Charles Bronfman Institute of Personalized Medicine. The BioM? Biobank is a prospective cohort study with over 40,000 ethnically diverse patients recruited from primary care and subspecialty clinics within the Mount Sinai Health System. The Institutional Review Board approved the study and informed consent was obtained for all subjects.

### Manual Review Study Design

In order to assess the accuracy of NAFLD patient selection, we performed a manual review on three different methods for patient selection. The study design is shown in Figure 1.

**Figure 1:**
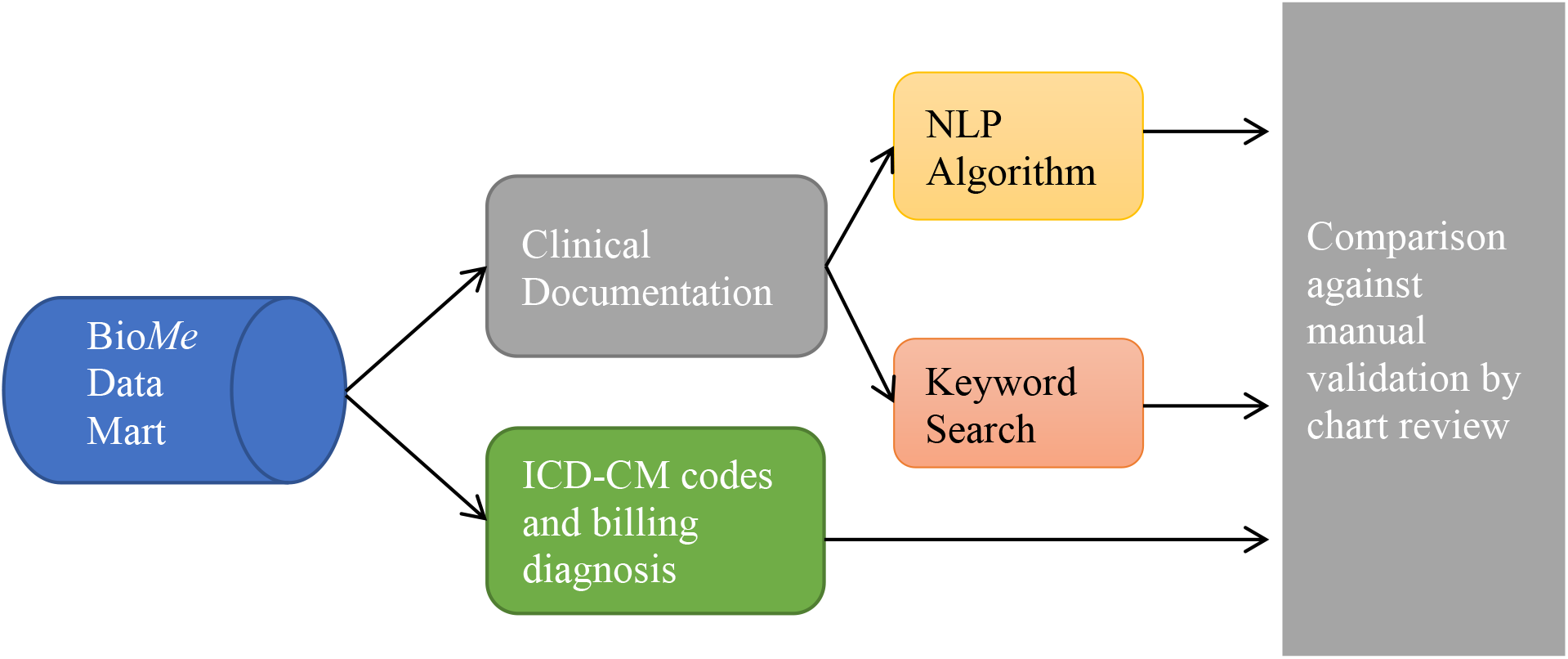
Study design for data extraction and manual chart review assessing automated identification of patients with NAFLD

We extracted the clinical documentation for all BioMe participants from the centralized DataMart up to December 31, 2017, with the sample starting June 11, 2009. Clinical documentation was comprised of progress notes, radiology reports, discharge summaries and pathology reports. After parsing the notes using NLP queries (described below), we conducted a simple text search for pre-defined phrases pertaining to NAFLD.

We compared both text-based approaches (NLP/text search) to manual validation using chart review in 100 randomly selected patients. We also estimated the prevalence of NAFLD in the BioMe cohort by each approach and compared them to prevalence estimates that have been reported in similar populations.[10] Finally, we explored the relationship between documentation on NAFL on an imaging test, contemporaneous acknowledgment of the possible diagnosis in a clinic note (suggesting that a responsible provider acknowledged the possible diagnosis and considered further evaluation or management), and subsequent documentation of NASH or cirrhosis in a provider note or discharge summary (potentially suggesting disease progression during the observation period).

### Approaches to Identify NAFLD in Electronic Health Records

#### Natural Language Processing

We used the CLiX clinical NLP engine produced by Clinithink (see Appendix for more information). CLiX is a general-purpose stochastic parser, which maps facts described in clinical narrative to post-coordinated SNOMED expressions, thereby creating a highly descriptive, standardized data layer capturing all identifiable clinical facts. SNOMED CT is a granular, hierarchical, general-purpose clinical terminology combining the most comprehensive single English terminology for medicine with a compositional grammar specifying how SNOMED CT concepts should be combined with expressions that define the clinical context around the concept. The combination of multiple SNOMED concepts to describe each identified patient fact provides infinitely more expressivity than possible with single concepts with post-coordination,[11] allowing each expression to include meta-data accounting for contextual differences affecting the meaning of the core SNOMED concepts identified. Supplementary Appendix Figure 1 demonstrates the SNOMED expressions identified for a sample phrase, and how the core concepts fit into the SNOMED CT terminology.

To identify patients matching specific criteria, we used a SNOMED query engine (a second component of CLiX) to perform hierarchical subsumption queries identifying relevant SNOMED expressions, meaning the identification of all SNOMED expressions found for the patient that were logical descendants (according to the SNOMED CT hierarchy) of each query expression. For example, according to SNOMED, *NAFLD* is a great-grandchild of *Disease of liver* and the temporal context of *Current or past* is a child of *Temporal context value.* The complete query for patients known to have had some form of liver disease is shown in Figure 2.

**Figure 2:**
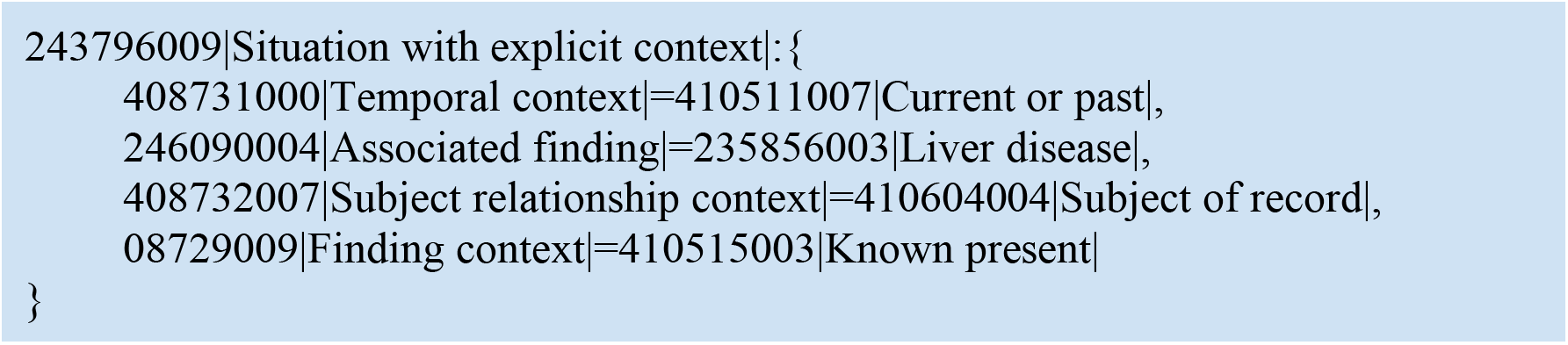
SNOMED expression/query for “disease of liver”

We used a SNOMED browser to identify other concepts critical to the identification of patients with NAFL that were not logical descendants of NAFL to ensure that we captured queries critical to the analysis. This led us to conclude that the optimal method would be to search for patients with the NAFL parent, *Steatosis of liver,* and exclude diagnoses other than NAFL. As a result, we excluded patients with Wilson disease, abetalipoproteinemia, alcoholic fatty liver, alcoholism (e.g. problem drinker, very heavy drinker, heavy drinker) and hepatitis B and C using a combination of NLP and ICD9/10 criteria.

#### ICD Codes and Simple Text Search to Identify NAFLD

We compared NLP-based approaches to other approaches for identifying NAFLD within the EHR. These included commonly used ICD9 and ICD10-CM codes for identifying disease. We also performed a text search (using SQL) to identify patients with notes containing phrases indicating NAFLD to see how well patients could be identified from notes without full NLP assessing negation, context, etc. ICD concepts selected included:

- ICD-9 571.8: Other chronic nonalcoholic liver disease
- ICD-10 K76.0: Fatty (change of) liver, not elsewhere classified
- ICD-10 K75.81: Nonalcoholic steatohepatitis (NASH)

Text search was performed for “NAFLD”, “NASH”, “fatty liver”, “steatosis”, “steatohepatitis”, “fatty infiltration of the liver” and “fatty infiltration of liver”.

#### Comparison with Manual Validation through Chart Abstraction

Two physicians independently, in a blinded manner, without knowing case/control status, reviewed all records for 100 patients identified as NAFLD and 100 patients identified as controls. Patients were classified as cases or controls based on clinical criteria. A clinical diagnosis of NAFLD required 1) the presence of fatty infiltration of the liver on imaging, 2) exclusion of hepatitis C infection, 3) absence of documented alcohol abuse. For controls we selected patients with a similar data density of imaging reports and progress notes but who did not fulfill the full NLP query. The two physician raters agreed in 95% of cases. Discordant cases were reviewed until consensus was achieved for all patients.

### Evolution and Progression Analyses of NAFLD

We used evolution analysis to examine how often patients identified as having NAFLD progressed to NASH or cirrhosis. We ordered narrative documents chronologically and then stepped through them, observing document type along with the presence of NAFLD and NASH/cirrhosis to observe how references moved between note types over time along with progression. From the first note on each patient, we stepped through looking for notes with a NAFLD reference and documented whether it was a radiology note or a clinic note (a provider note or a discharge summary). Once a radiology note had been identified, we tracked how long it took for a non-radiology clinical note to reference NAFLD. If no further non-radiology notes mentioned it, we confirmed that there were additional clinical notes where NAFLD was not mentioned to ensure that lack of further reference was not due to a discontinuation of care. Finally, the evolution analysis was combined with the progression analysis, and we examined patients where NAFL was identified by radiology, but was not recognized within the clinical documentation and yet the patient developed NASH or cirrhosis.

### Statistical Analyses

We calculated summary statistics to determine precision, recall, F1 (a measure of test accuracy that considers precision and recall) and F2 scores (placing more emphasis on recall and thus emphasizing false negatives more than false positives).[12] We assessed ICD-9/10 codes, NLP, and text search for their ability to accurately identify NAFLD patients against the reference-standard (manual abstraction), calculating precision, recall, false positive rate, F1 and F2 scores for each method. We compared estimates of F1/F2-scores between algorithms using the McNemar test. We used generalized score statistics to compare precision, recall and the false positive rate. All data analysis was performed in Python (with standard packages). The significance threshold for analyses of differences was calculated as a two-sided significance p-value of <0.05.

## RESULTS

We included 7,766,654 notes of 38,575 Bio*Me* enrollees from July 8, 2002 through December 31, 2017. Parsing with NLP yielded 428,469,717 post-coordinated SNOMED expressions describing clinical concepts and related context. Figure 3 shows the queries (or query clusters, in the case of alcohol users) for the identification of case and control patient cohorts.

**Figure 3:**
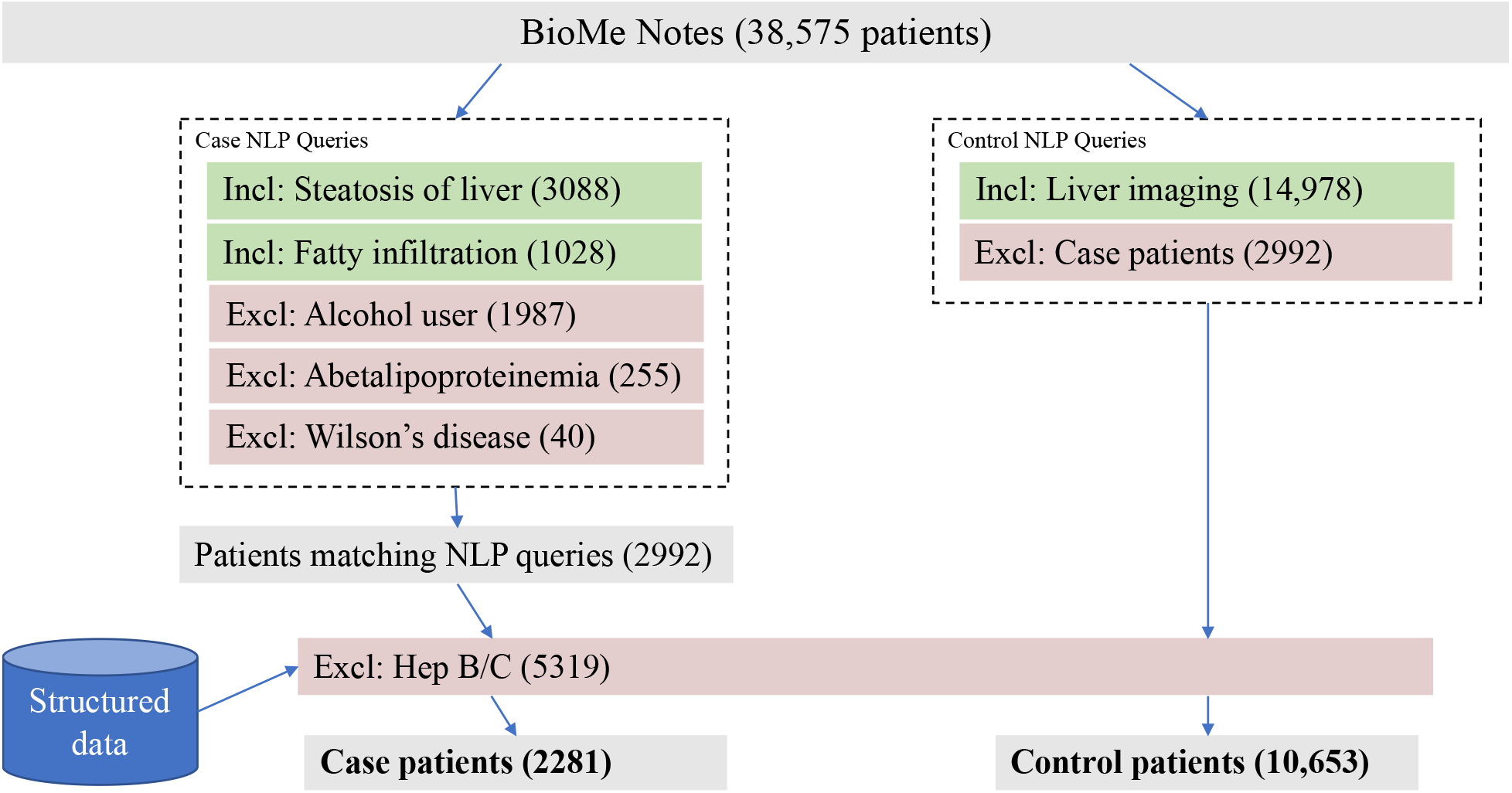
NLP SNOMED queries for case and control cohorts

### Baseline characteristics for NAFLD case and control cohorts

As the NLP-based approach had the best overall summary statistics, we used the cases and controls identified by this approach for further analyses. The baseline characteristics of the 2281 cases and 10,653 control participants are shown in Table 1.

**Table 1:** Baseline Characteristics of NAFLD Cases and Controls

|  | Cases (n=2281) | Controls (n=10,653) | P |
| --- | --- | --- | --- |
| Mean Age | 59.8 | 59.5 | 0.01 |
| % Male | 42% | 39% | 0.01 |
| *Race* |  |  | <0.01 |
| African American | 20% | 30% |  |
| Caucasian/European | 23% | 22% |  |
| Asian | 2% | 2% |  |
| Hispanic | 48% | 41% |  |
| Other | 6% | 5% |  |
| *Mean liver serology at baseline* |  |  |  |
| Aspartate Aminotransferase | 45.7 | 39.7 | 0.01 |
| Alanine Aminotransferase | 48.5 | 35.9 | <0.01 |
| *Baseline Comorbidities* |  |  |  |
| Diabetes Mellitus, n (%) | 1233 (40.7%) | 3524 (27.6%) | <0.01 |
| Hypertension, n (%) | 2079 (68.6%) | 7898 (61.9%) | 0.01 |
| Mean Body Mass Index | 31.8 | 29.2 | <0.01 |

### Identification of NAFL by different approaches

We identified 2281 cases of NAFL using NLP and 10,653 patients appropriate as controls for manual review. We also identified 1232 patients by ICD codes and 5489 patients by text search. The overlap patients identified by different approaches are shown in Figure 4.

**Figure 4:**
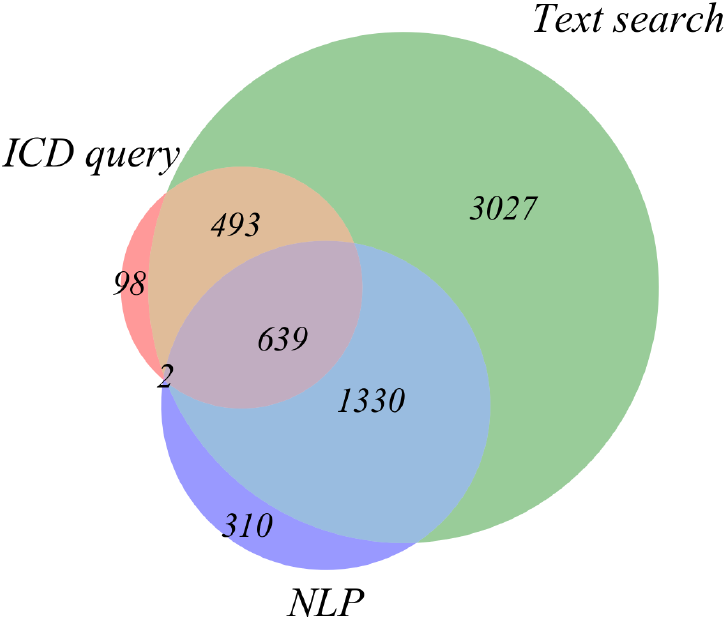
Overlap of patients identified by three different approaches

#### Comparison between different approaches

The summary statistics for a manual validation comparing the three patient selection methodologies on 100 cases and 100 controls are shown in Table 2.

**Table 2:** Accuracy of NLP identification of NAFLD patients relative to ICD codes and basic text search

|  | **NLP** | **ICD Selection** | **Text Search** |
| --- | --- | --- | --- |
| *Precision/PPV* | 0.89 | 0.97 | 0.83 |
| *Recall/Sensitivity* | 0.93 | 0.28 | 0.81 |
| *Specificity* | 0.89 | 0.99 | 0.84 |
| *FPR* | 0.11 | 0.03 | 0.17 |
| *F1* | 0.91 | 0.44 | 0.82 |
| *F2* | 0.92 | 0.34 | 0.81 |

Overall, the NLP approach had the best summary statistics with high precision (0.89), recall (0.93) and F1 scores (0.91) and low FPR (0.11). Precision of the ICD approach was higher but recall and the F1/F2 scores were significantly lower. The text-based approach was significantly worse than the NLP approach with respect to all parameters (p<0.05).

#### Reasons for misidentification of NAFL

We analyzed reasons for false positives and false negatives of the NLP algorithm. Most false positives were due to hypothetical or otherwise uncertain references. Three examples are shown in Table 3.

**Table 3:** Review of NLP false-positives

| **Problem** | **Example** |
| --- | --- |
| Hypothetical reference | Nonspecific hepatic parenchymal change, such as, but not limited to, fatty liver |
| Hypothetical reference | It is likely that his ongoing moderate alcohol use and NAFLD (nonalcoholic fatty liver disease) are contributing to this elevation |
| Uncertain reference | Liver is nodular in contour, suggestive of cirrhosis. There is diffuse hypoattenuation of the liver likely representing hepatic steatosis |

The low precision and F1 score of the ICD-based approach occurred because ICD codes are more specific than sensitive. An ICD code is only applied if the physician and hospital coders are certain that the condition exists or for billing purposes, but there are many reasons (often nonmedical) that a problem might not be coded despite its presence. Text search had a high false-positive rate due to negations, references in templates and other references not indicative of the patient having the problem. The major limitations of each approach are explained in Table 4.

**Table 4:** Strengths and limitations of patient selection modalities

| **Methodology** | **Strengths** | **Weaknesses** |
| --- | --- | --- |
| *ICD query* | Very high specificity<br>Can be standardly used | Low sensitivity |
| *Text search* | Moderately high sensitivity and specificity | Unpredictable as it depends on identifying the exact phrases used by note authors<br>Scalability issues since manual verification of exact search strings to be used |
| *NLP* | High sensitivity and specificity | Requires access to NLP infrastructure, especially capable of post-coordinating SNOMED expressions |

### Progression from NAFL to NASH/Cirrhosis

Among 2281 patients identified as having NAFL, 486 (21.3%) were identified as having NASH. Another 187 were identified as developing cirrhosis, not specifically due to NAFL. Among the 602 documented as having both NAFL and NASH, 310 patients had NAFL and NASH identified at the same time. For the remaining 176 where NAFL was identified prior to NASH, the median progression time to documentation of NASH was 410 days.

### Evolution Analyses of Information Flow from Radiology to Clinical Notes

Of the 2281 patients with NAFLD identified in notes, 619 had NAFLD identified in only radiology notes (excluding pathology), 1020 had it identified in only clinical documentation and 619 again identified it in both radiology and clinical notes. A small number of patients (23) had NAFLD identified only in pathology notes, but these references were excluded from all analyses. (Figure 5)

**Figure 5:**
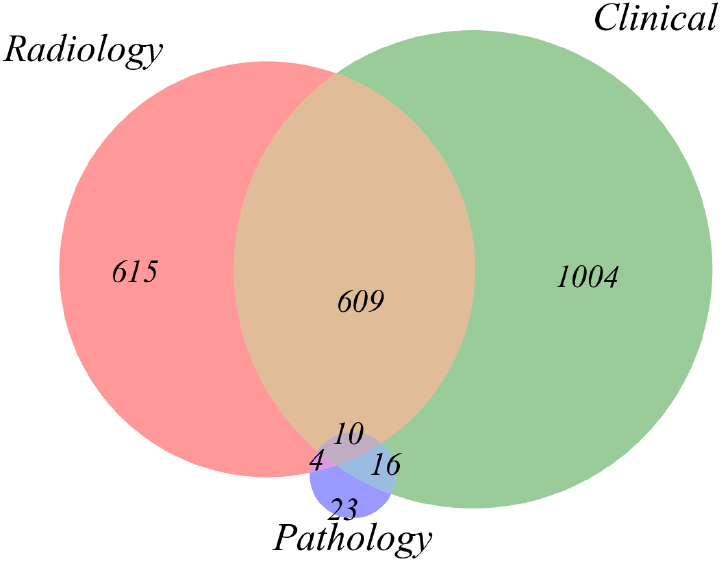
Distribution of note types referencing NAFLD, counted by patient with at least one note of each type referencing NAFLD

Of the 619 patients with NAFL (fatty infiltration/steatosis) identified in radiology notes and not acknowledged in clinical documentation, 232 (37.5%) developed NASH or cirrhosis. To ensure this did not include clinicians immediately reacting to the NAFLD identified by radiology, we omitted patients where NASH was identified on the same day as the radiology notes identifying NAFLD, which brought the total to 170 (27.5%). After excluding same-day identification, we observed a temporal gap averaging 1057.3 days (range 4 to 4324 days). Of the 170 patients identified, 105 had NASH and the remaining 65 had cirrhosis.

## DISCUSSION

We assessed several informatics approaches to identify NAFLD within the EHR data compared to manual validation by clinicians in a large, multiethnic cohort. Our observations suggest that NLP approaches had the best overall performance compared to ICD and text search-based approaches. In addition, the prevalence of NAFLD (~18% in those patients with imaging data) identified by the NLP-based approach was similar to population prevalence using nationally representative data, especially considering the ethnic minority predominant demographics of the BioMe Biobank.[2, 10]

The widespread availability of EHRs in hospital systems provides an opportunity for clinical and genomic research, population health analytics, and improvement of patient care through clinical decision support.[13, 14] However appropriate use of the large-scale, granular information in EHRs depends on accurate and rapid identification of patients with the disease of interest. This is especially relevant to the study of NAFLD where the study of its natural history has been restricted by the lack of large, longitudinal cohorts with most cohorts comprising a few hundred radiological/histologically confirmed NAFLD patients.[15, 16] High-throughput identification of NAFLD with “electronic” follow-up through the EHR, could aide in understanding the risk factors for progression to cirrhosis.

Previous studies have attempted to create algorithms to identify NAFLD through the EHR. A study by Corey et al. used limited natural language processing approaches to define NAFLD within the EHR confirmed through radiology reports in combination with ICD codes.[17] They demonstrated that the PPV (89%) and NPV (56%) was superior to an approach utilizing ICD-9 coding alone or a model incorporating AST/ALT laboratory values. However, the language parsing approach used counted only the occurrences of pre-defined terms related to NAFLD without considering critical issues in NLP including negation, context, spelling, and acronyms.[18, 19] In contrast, we used an integrated NLP approach that fully accounted for these issues as applied to EHR documentation. On manual validation, we demonstrated improved summary statistics compared to not only ICD-9/10 codes, but also to simple text search.

While conducting analyses of the information flow between several different types of documentation where NAFLD may be identified, we found that NAFL discovered in radiology notes was not acknowledged in clinical documentation in approximately one-half of the cases. One in ten patients who had NAFL identified in radiology notes but which was not acknowledged in clinical documentation were later documented as having NASH without any acknowledgement of NAFL in the intervening clinical encounters or progress notes.

Previous studies have demonstrated that findings on radiology reports (particularly incidental findings) are not uniformly acknowledged or pursued by relevant providers.[20, 21] This may in part reflect the deluge of biomedical information available within EHR systems, which may lead to a breakdown in information flow.[22] Although, progress has been made on “closing the loop” on these non-emergent radiology findings,[23] it still is an active area of both research and clinical improvement.

Past studies have demonstrated that NLP can be used to obtain valuable data for research that can be more accurate than ICD codes.[7,24–26] This study supports these findings, identifying NLP as clearly superior for individual phenotype algorithms. As data volume and accuracy are critical for big data initiatives, it stands to reason that NLP-derived features will yield superior models for these endeavors.

Our research should be interpreted in the context of its limitations. First, we used only one NLP tool for analyses. This approach could be accomplished with any NLP software capable of accurately mapping to and querying any clinical terminology as expressive as SNOMED CT/DL, however no other tool capable of this task was available at the time of publication.

Second, we used data from only one medical center thus the applicability of our findings to other settings remains to be determined. That said, the Mount Sinai medical system has a large network of providers from different specialties and each with their own unique writing style, therefore we believe this electronic phenotyping approach can be successfully used across multiple healthcare systems.[27]

Third, in the analyses of information flow from radiology to clinical documentation, it is possible that patients may have been receiving care outside of our hospital system and thus not be captured by our EHR. However, we limited this possibility by conducting a subset in which patients had at-least one clinical encounter after the radiology identification of NAFLD.

In summary, we demonstrate that NLP-based approaches have superior accuracy in identifying NAFLD within the EHR compared to ICD/text search-based approaches. There is lack of acknowledgement in clinical documentation of NAFL findings in radiology reports and a significant number of these patients are later reported to have NASH. Our observations suggest that NLP-based approaches have the potential to identify clinically relevant observations in EHR data that warrant additional follow-up.

## ACKNOWLEDGEMENTS

The Bio*Me* healthcare delivery cohort at Mount Sinai was established and maintained with a generous gift from the Andrea and Charles Bronfman Philanthropies. This work was supported in part through the computational resources and staff expertise provided by the Department of Scientific Computing at the Icahn School of Medicine at Mount Sinai.

## Funding

SGC is supported by the National Institute of Diabetes and Digestive and Kidney Diseases (NIDDK) (grant no. R01DK096549 to SGC). GNN is supported by a career development award from the National Institutes of Health (NIH) (K23DK107908) and is also supported by R01DK108803, U01HG007278, U01HG009610, and 1U01DK116100-01 grants. SGC and GNN are members and are supported in part by the Chronic Kidney Disease Biomarker Consortium (U01DK106962). SGC is also supported by R01DK106085, R01HL85757, R01DK112258, and U01OH011326 grants. LC is supported in part by the NIH (5T32DK007757 – 18) and the recipient of a research grant from the Renal Research Institute.

## Conflicts of interest

TTVV was part of launching Clinithink and retains a financial interest in the company. GNN is cofounder of Renalytix AI and owns equity in that company.

## APPENDIX

Our NLP system, CLiX, is produced by Clinithink (clinithink.com), a clinical NLP and analytics company based in the UK. CLiX represents all identified patient facts using post-coordinated SNOMED expressions. Supplementary Figure 1 demonstrates encoding for two phrases: “fatty liver” and “no alcohol abuse”. The diagram to the right shows how the concept for “fatty liver” (Steatosis of liver) fits into the SNOMED hierarchy.

**Supplementary Figure 1:**
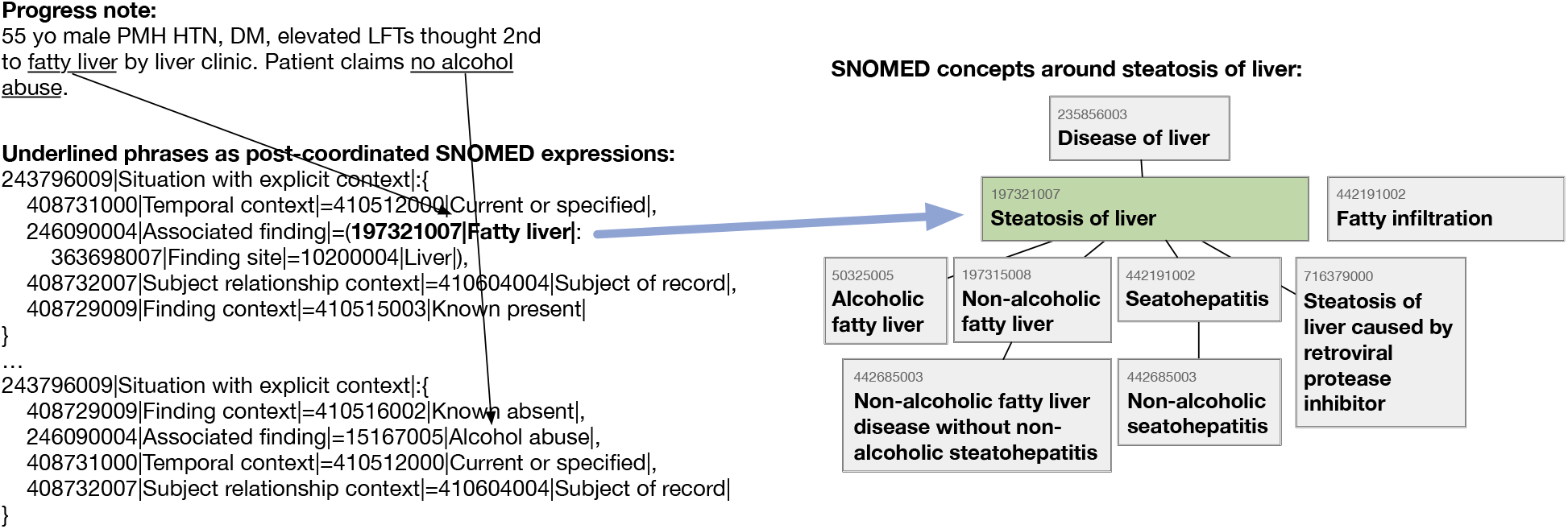
Sample CNLP encoding

